# Scene Structure Predicts Perceptual Decisions in Naturalistic Detection Tasks

**DOI:** 10.64898/2026.06.03.729800

**Authors:** Jun Yang, Tiziana Vercillo, Teresa Emma Cutrona, Simone Azeglio, Giandomenico Iannetti, Peter Neri

## Abstract

The human visual system can identify objects in complex natural scenes, yet the mechanisms supporting robust perception under such variable conditions remain incompletely understood. Here, we investigate how the statistical structure of natural scenes shapes perceptual evidence formation and determines whether near-threshold stimuli are perceived correctly or incorrectly. We combine controlled psychophysics, outdoor augmented reality (AR), deep neural networks (DNNs), image-feature analysis, and EEG to examine how background context modulates perceptual decisions. Across multiple detection tasks, human performance was systematically influenced by probe-free background structure. DNNs trained on background images alone predicted correct and incorrect behavioral outcomes, with stronger effects in postcue conditions, suggesting that global scene context contributes to local perceptual decisions when spatial uncertainty is higher. AR experiments further showed that these context-driven effects persist in naturalistic viewing environments. To identify the visual information underlying these effects, we analyzed low-level image statistics. Texture entropy and edge density emerged as informative features, and conventional machine-learning models trained on these measures achieved meaningful correct/incorrect classification. Finally, EEG analyses revealed neural signatures of image-driven perceptual variability: activity during the probe-free preimage window distinguished later correct from incorrect trials, and combining EEG with image-derived features improved decoding performance. Together, these findings show that perception in natural scenes is not determined solely by the target, but is shaped by the statistical structure of the surrounding context. By linking psychophysics, AR, DNN modeling, image statistics, and EEG, this work provides a unified framework for understanding how environmental structure and neural dynamics jointly support perceptual decision-making.

## 1 Introduction

Human vision operates in environments filled with complex textures, surfaces, and spatial structure. Although visual detection is often studied by manipulating local target properties, such as contrast, size, or eccentricity, perceptual decisions in natural scenes rarely depend on the target alone. The same faint probe can be detected easily in one background but missed in another, suggesting that local perceptual decisions are strongly constrained by the global structure of the surrounding scene. Understanding how global scene statistics shape perceptual success or failure is therefore essential for explaining vision under naturalistic conditions.

Natural scenes contain rich and reproducible statistical regularities, including luminance distributions, edge structure, texture complexity, spatial correlations, and surface organization. These regularities are not merely visual noise; they provide the statistical context within which local stimuli are encoded and interpreted. For example, backgrounds with dense edges, high texture entropy, or abrupt luminance transitions may interfere with the detection of weak local targets, whereas simpler and more homogeneous regions may facilitate detection. Thus, perceptual errors should not necessarily be treated as random lapses, but may instead reflect systematic interactions between local targets and global scene structure.

Previous studies have shown that natural scene statistics influence visual sensitivity, attention, masking, and recognition. However, it remains unclear whether the global structure of a background alone is sufficient to predict trial-by-trial perceptual outcomes. In other words, if the target itself is removed from the image, can the remaining background still indicate whether a human observer is likely to respond correctly or incorrectly? Answering this question would provide direct evidence that perceptual variability is partly determined by scene-level structure, rather than being driven only by target properties or internal decision noise.

Here, we address this question by combining behavioral experiments, augmented reality, deep neural network modeling, and EEG analysis. Human observers performed visual detection tasks in which faint probes were embedded in natural images or real-world augmented-reality environments. Across trials, probe properties were controlled, while the surrounding scene structure varied. Behavioral responses were then categorized as correct or incorrect, allowing us to examine whether different backgrounds systematically led to different perceptual outcomes.

A key aspect of our computational analysis is that the models were trained on probe-free images. As shown in Fig. 1, human observers viewed images that could contain a faint probe and made detection responses. In contrast, the deep neural networks received only the background image after the probe had been removed, and were trained to predict whether the corresponding human response was correct or incorrect. This design forces the model to rely exclusively on probe-free background context. Therefore, above-chance prediction would indicate that the background itself contains sufficient statistical information to bias human perceptual decisions.

**Figure 1:**
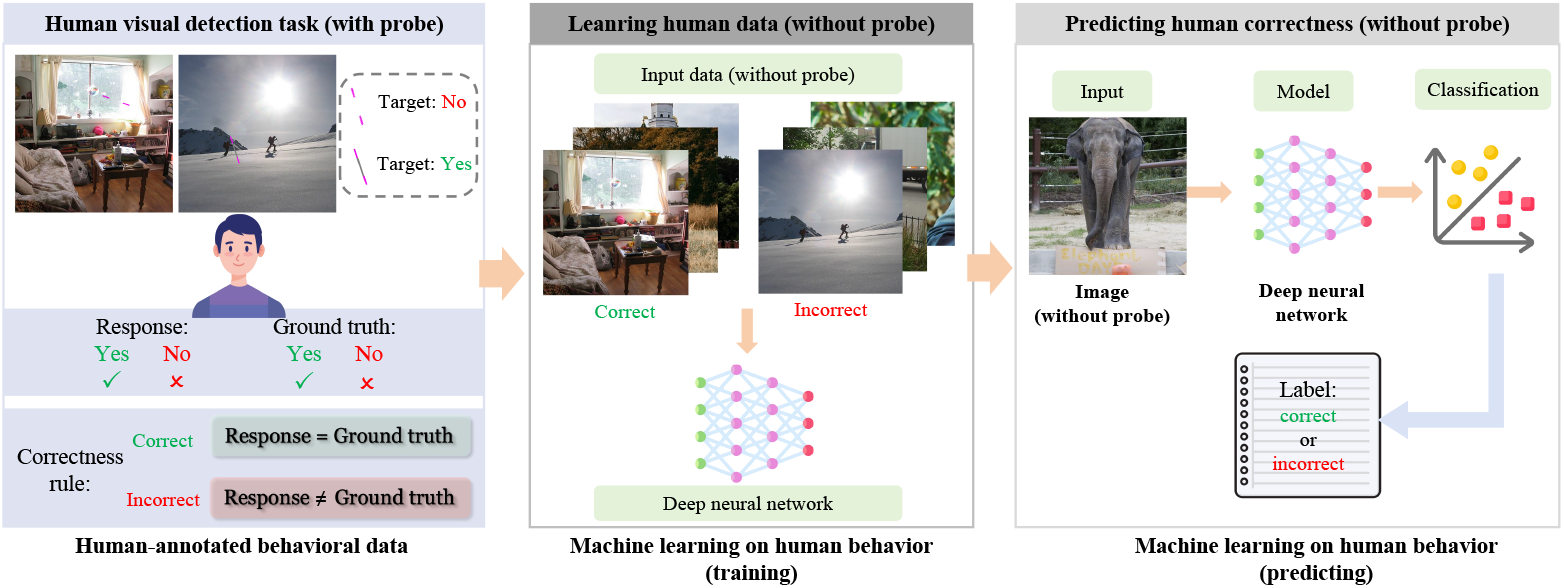
Relationship between the human detection task and the machine-learning analysis. Human observers performed a detection task on images that could contain a faint probe, producing correct or incorrect responses. For machine learning, the probe was removed, and deep neural networks were trained to predict the human perceptual outcome from the background image alone. This probe-free design tests whether global scene structure is sufficient to predict human perceptual decisions.

We further extended this approach from conventional screen-based psychophysics to augmented reality. In the screen-based experiments, each trial presented a different natural image, allowing us to measure how trial-by-trial changes in global image structure affected detection performance under controlled conditions. In the augmented-reality experiments, virtual probes were embedded into stable real-world environments, preserving spatial continuity, depth cues, lighting, and motion parallax. This allowed us to test whether context-dependent detection effects also occur under more naturalistic viewing conditions. Across both settings, we ask whether local detection judgments are systematically shaped by the structure of the surrounding visual environment.

Finally, we examined whether these scene-driven effects are reflected in neural activity before the target appears. EEG was recorded while participants performed the detection task, and we focused on a pre-probe time window in which the background scene was visible but the probe had not yet been presented. If EEG activity during this window predicts whether the subsequent response will be correct or incorrect, this would suggest that the brain’s response to the global scene already carries information about the upcoming perceptual decision. In this way, EEG provides a neural counterpart to the image-based computational analysis.

Together, these components provide a multimodal account of how background context constrains local perceptual decisions. Behavioral experiments test whether detection performance varies systematically across scenes; deep neural networks trained on probe-free images assess whether background structure is computationally sufficient to predict human outcomes; augmented-reality experiments examine whether these effects generalize to naturalistic viewing; and EEG analysis evaluates whether scene-driven biases are reflected in neural activity before probe onset.

In summary, this paper investigates whether the statistical structure of natural scenes can predict whether a local target will be seen or missed. We show that perceptual outcomes are not determined solely by the target itself, but are systematically modulated by the surrounding visual context. By combining psychophysics, augmented reality, deep neural networks, and EEG, this work provides an ecologically grounded account of visual detection in which background structure plays a central role in shaping local perceptual decisions.

## 2 Human visual perception in online experiments

### 2.1 Dataset

The natural images are from two datasets, COCO dataset [1] and SAM dataset [2]. The COCO dataset [1] is a large-scale benchmark for object detection, segmentation, and scene understanding. It contains over 200,000 images with more than 80 object categories annotated with bounding boxes, instance segmentation masks, and keypoints. Unlike earlier datasets that focus on iconic object-centric images, COCO emphasizes objects embedded in complex everyday scenes, thereby promoting models that capture contextual information and robust visual representations. The SAM dataset[2] is a large-scale image segmentation dataset introduced alongside the Segment Anything Model (SAM). It contains over 11 million images and more than 1 billion high-quality segmentation masks, making it one of the largest segmentation datasets to date.

### 2.2 Segment experiment

#### 2.2.1 Experiment setting

The stimuli used in this experiment consisted of natural images along with their corresponding cues and probes. The natural images were sampled from the COCO dataset [1]. We selected images whose width or height exceeded 480 pixels and center-cropped them to a resolution of 480 × 480 pixels to ensure sufficient spatial detail for the detection task.

The cues were constructed from two magenta line segments, each measuring 5 pixels in width and 29 pixels in height. The probe was also a magenta segment of the same dimensions. The cues and probe were generated in matched pairs, and their positions were randomly placed within an annulus-shaped region: they were constrained to be outside a 15% border margin and outside a central rectangle spanning ± 5% around the image center.

The experimental design is illustrated in Fig. 2, which includes two conditions: precue and postcue. Each natural image was shown twice per session—once in each condition. Across trials, natural images were divided evenly into probe-present and probe-absent conditions. In probe-present trials, the probe was embedded at a location between the cue segments; in probe-absent trials, the images remained unaltered.

**Figure 2:**
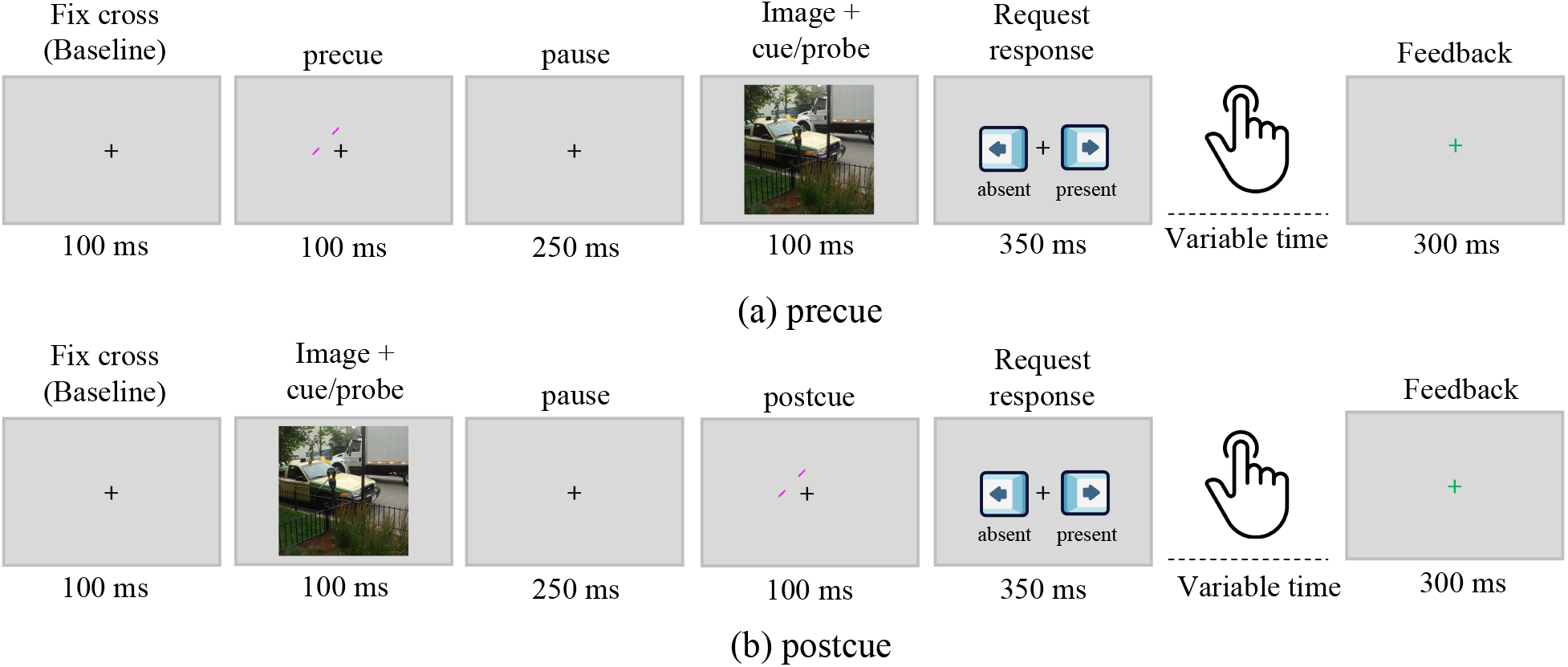
The pipeline for precue and postcue conditions. In the precue condition, a cue is presented first, followed by a brief blank interval, after which the natural image (with or without a probe) appears. In this condition, participants know the potential probe location in advance and can rely primarily on local information. In the postcue condition, the order is reversed: participants first view the natural image, then—after a blank interval—receive the cue. Because the natural image precedes the cue, participants must rely more heavily on global scene information to detect the probe.

We manipulate cue timing using precue and postcue conditions. In the precue condition, participants saw the cue first, so they knew where to attend before the image appeared. This means they could focus more on the local region around the target location. In the postcue condition, participants first saw the whole image and only later received the cue. Therefore, they had to process the global image structure before knowing the exact target location. In the precue condition, the cue was displayed for 100 ms, followed by a 250-ms blank screen, and then the natural image (with or without a probe) was presented for 100 ms. In the postcue condition, this order was reversed. The natural image appeared for 100 ms, followed by a 250-ms blank interval, and then the cue was shown for 100 ms. Given the 250-ms interval, a visual masking effect is unlikely to occur [3, 4], ensuring that the cue in the postcue condition does not provide probe-location information. Thus, probe location is available only in the precue condition. The precue and postcue conditions are mixed in each block. By comparing precue and postcue conditions, we can test whether global scene information plays a stronger role when spatial uncertainty is higher.

#### 2.2.2 Participants and data preparation

We conducted the experiment on Prolific with data collected from 52 subjects/sessions. Each participant completed 1,000 precue and 1,000 postcue trials presented in a randomized order. In total, we collected 51,413 usable trials across the precue and postcue sessions. The mean accuracy in the postcue condition was 71.37%, whereas the accuracy in the precue condition was 78.16%. This difference is expected: because the cue precedes the image in the precue condition, participants can use precise local information, making the task easier.

#### 2.2.3 Results

We have 13,724 Correct/Incorrect trials. The AUC results for precue and postcue conditions across models are shown in Fig. 3. The first 4 groups represent the performance of different DNN architectures: ResNet [5], EfficientNet [6], ConvNeXt (tiny) [7], and Swin Transformer [8]. The last boxplot is a control experiment, showing the AUC is about chance when shuffling the labels.

**Figure 3:**
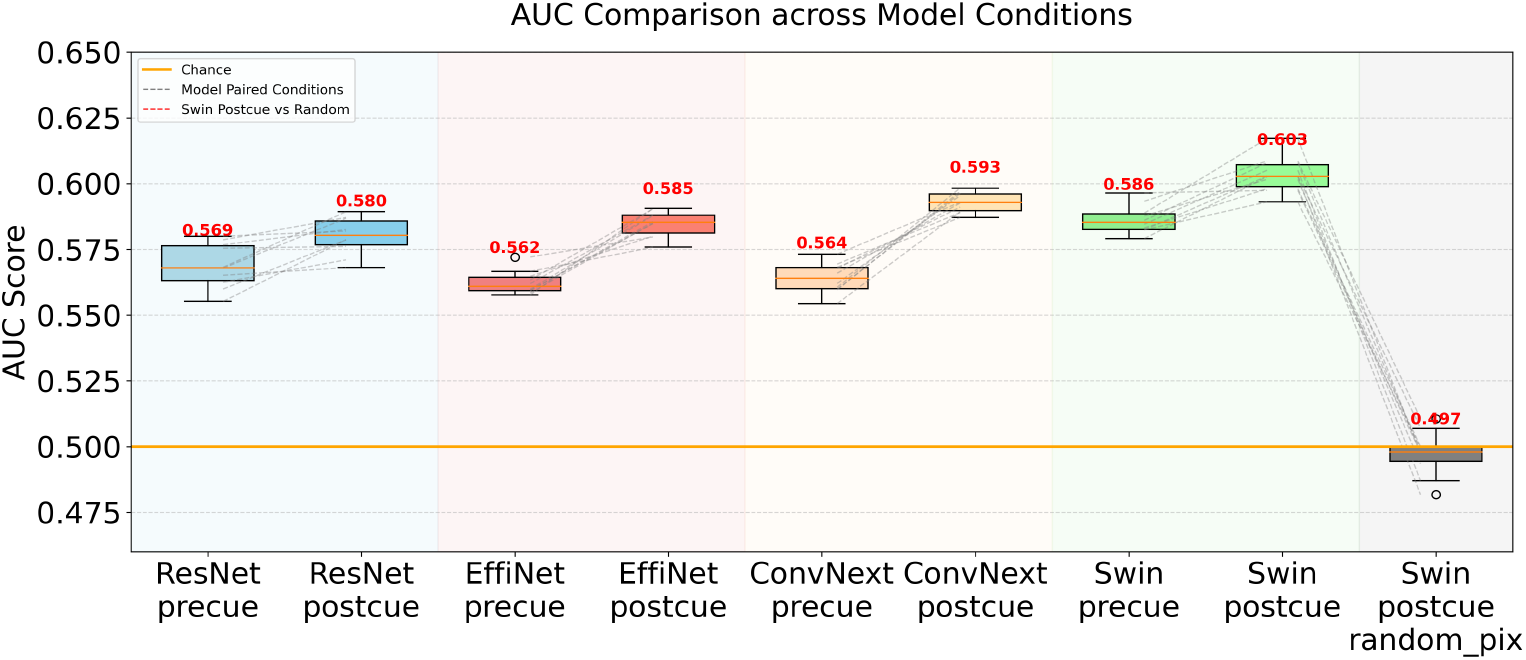
AUC comparison across models for RGB images. Different background colors correspond to different DNN architectures. The grey dashed line indicates trial-by-trial comparison.

There are two main findings showing in this figure:

- Model’s prediction AUCs are all above chance.
- Prediction AUC of postcue condition is higher than precue.

The first finding indicates that clean background image alone is sufficient to classify Correct versus Incorrect trials, meaning scene alone can modulate local perceptual processes independently of the target itself. The second finding shows human rely on more global information. Comparing precue and postcue results, the postcue condition consistently achieves higher AUC values. This difference is theoretically meaningful: precue trials encourage the use of local information, whereas postcue trials require integrating global information to locate the probe. Since global scene context influences local feature detection—as demonstrated in prior work [9]—the higher AUC in postcue suggests that global information is a stronger predictor of human performance. In other words, the global scene impacts how does local perception work.

### 2.3 Blob experiment

In the previous task, the probe was a grey bar that could appear at any position and freely rotate from 0^*°*^ to 360^*°*^. As illustrated in Fig. 4, when the orientation of the probe happened to coincide with edges or elongated structures in the scene—such as benches, railings, or vehicles—the probe could become perceptually fused with the background, thereby reducing its visibility. Because the probe is an oriented bar, it can incidentally align with object contours, allowing it to blend into the image structure.

**Figure 4:**
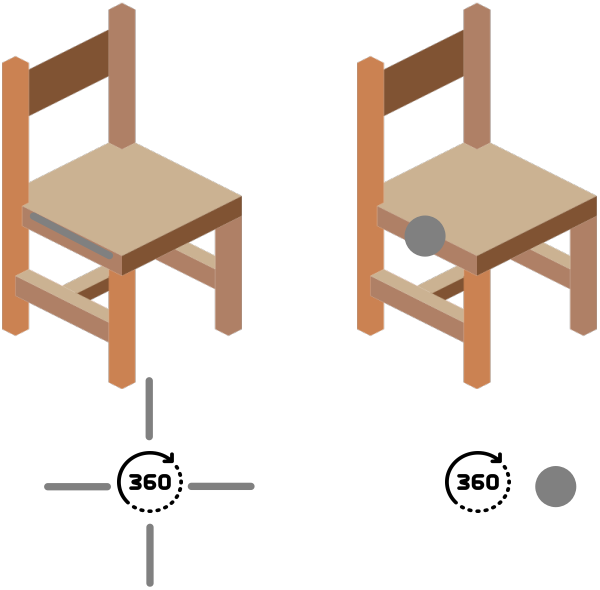
Comparison between bar (left) and blob (right) stimuli. Because the bar probe can appear anywhere and rotate from 0^*°*^ to 360^*°*^, it sometimes aligns with existing object edges in the scene, making it difficult to detect.

To examine whether the classification effect depends on such orientation-specific cortical processing, we conducted a follow-up experiment in which the probe shape was changed from a grey bar to a grey disk. This manipulation removes the orientation dimension entirely, allowing us to test whether the classification effects persist when orientation-related neural activation is eliminated.

#### 2.3.1 Experiment setting

In this new experiment, the probe was a grey blob instead of a bar, and the cue was redesigned as a magenta circle. All other experimental parameters were identical to the previous experiment. By replacing the oriented probe with a symmetric blob, the orientation variable was eliminated, enabling us to test whether the classification effect generalizes beyond orientation-driven cortical mechanisms.

#### 2.3.2 Participants and data preparation

A total of 50,410 trials were collected for the blob experiment. Unlike the previous bar-based experiments—in which participants showed higher performance in the precue than the postcue condition—the average accuracy across participants was the same, i.e., 77.65% for both precue and postcue. This indicates that prior location information did not facilitate detection performance when the probe lacked orientation.

#### 2.3.3 Results

We selected 10,018 Correct/Incorrect trials as the training dataset and 1,250 trials as the testing dataset. For computational modeling, we trained three DNN architectures to classify these human-derived labels: ResNet [5], ConvNeXt (tiny) [7], and Swin Transformer [8]. The resulting AUC values are shown in Fig. 5.

**Figure 5:**
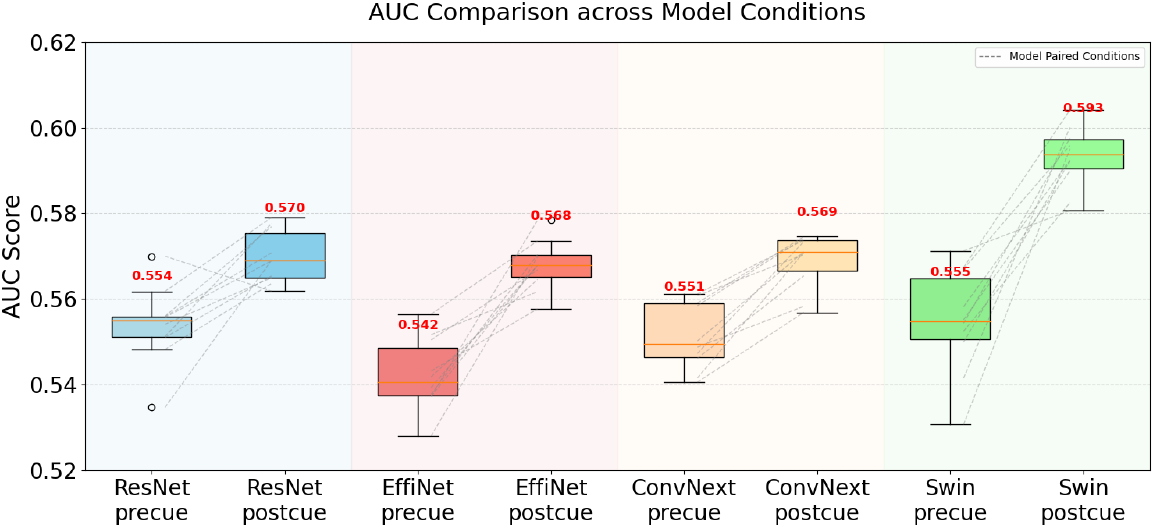
AUC comparison across models in the blob-based experiment. Although human detection accuracy was identical for the precue and postcue conditions (both 77.65%), all three models consistently achieved higher AUC values in the postcue condition than in the precue condition.

The results reveal three key findings. First, the classification effect persisted in the blob condition, demonstrating that the model can still predict human detection performance even when the probe carries no orientation information. Second, consistent with the earlier experiments, AUC values for the postcue condition were consistently higher than those for the precue condition across all three models. Third, this AUC advantage is not explained by differences in subject performance: even when participants showed comparable accuracy in precue and postcue trials, the classifier still achieved higher AUC in the postcue condition.

Crucially, this postcue advantage in model prediction emerged despite the fact that human behavioral accuracy did not differ between precue and postcue conditions. In other words, although participants performed equally well in the two cueing conditions, the models consistently achieved higher AUCs for postcue trials.

This dissociation argues against interpreting the segment-related classification effect as a trivial consequence of differences in human detection performance between cueing conditions. Instead, it suggests that DNNs exploit global contextual properties of natural images that correlate with perceptual difficulty in a way that is not directly reflected in overt accuracy measures.

Overall, these findings demonstrate that the observed classification effect is not driven by orientation-based cortical processing nor by differences in behavioral performance across cueing conditions. Rather, it reflects more general contextual properties of natural scenes that influence human detection performance, reinforcing the idea that global scene structure plays a central role in shaping visual perception.

Interestingly, although participants’ behavioral accuracy did not differ between precue and postcue conditions, the models nevertheless showed superior classification performance for postcue trials. This indicates that DNNs may continue to exploit global contextual features of the natural images—features that correlate with perceptual difficulty and remain informative even when orientation cues are absent.

Overall, these findings demonstrate that the observed classification effect is not driven by orientation-based cortical processing. Instead, it reflects more general contextual properties of natural images that influence human detection performance, reinforcing the idea that global scene structure plays a central role in shaping visual perception.

### 2.4 Understanding the experiment

DNNs were trained to predict these human-derived labels using only the original, probe-free images as input. Because the probe was never present in the model’s input, the network could not learn any direct mapping between the physical probe signal and behavioral outcome. Nevertheless, the models achieved classification performance reliably above chance, indicating that systematic properties of the background scenes are correlated with human success or failure in detecting weak signals. Crucially, when operating on probe-free images, the DNN does not encode probe-related information explicitly. Instead, it learns an *indirect* association mediated by human behavior. Human observers viewed both the probe and the surrounding scene, and their detection decisions were jointly influenced by the weak probe signal and the contextual properties of the image. As a result, the Correct/Incorrect labels implicitly embed a mixture of probe-related evidence and background-driven perceptual factors. Although the probe itself is absent from the input, the background scene retains contextual features that systematically biased human judgments, leaving a statistical trace in the labels that the DNN can exploit.

A comparison between postcue and precue conditions in both segment task and blob task further clarifies the role of global scene information. When putting all data together, across models and tasks, AUC is consistently higher in the postcue condition than in the precue condition (Fig. 6). In the postcue condition, observers must first encode the scene globally before receiving spatial information about the probe location, whereas the precue condition allows attention to be narrowly focused in advance. The higher predictability in postcue therefore indicates that global scene structure plays a more prominent role when detection relies less on anticipatory spatial attention and more on scene-level visual information.

**Figure 6:**
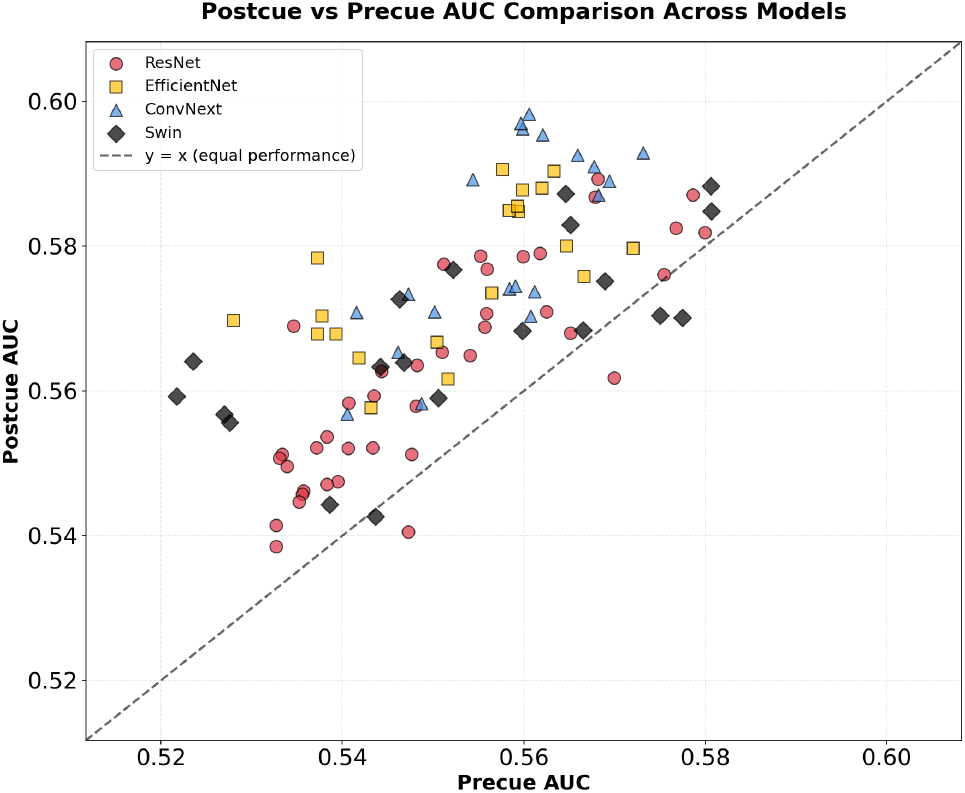
Postcue vs precue AUC comparison across models and tasks.

Taken together, these findings demonstrate that intrinsic visual properties of natural scenes systematically influence human sensitivity to faint signals, even when those signals are physically identical across trials. Human judgments are shaped not only by the presence or absence of the probe, but also by global scene context, which can facilitate or hinder detection. The ability of DNNs to predict human performance from background images alone reveals that scene structure contains information predictive of perceptual accuracy. This framework offers a new perspective on visual processing in complex environments and highlights the value of machine learning models as analytical tools for probing the contextual factors that shape human visual behavior.

## 3 Human visual perception in augmented reality experiments

We turn to augmented reality (AR) as a platform for studying visual detection under more naturalistic conditions. Classical visual psychophysics has typically been carried out in tightly controlled laboratory environments, where participants’ heads are immobilized, their gaze is constrained, and their behavioural repertoire is severely limited [10, 9]. More recent work has begun to relax these constraints using immersive virtual reality (VR), in which participants are free to move their head and body while performing visual tasks [11, 12]. These VR approaches are a major advance but still rely on fully simulated environments, which can differ substantially from the complexity and variability of real-world scenes.

AR provides an even closer approximation to natural visual experience. Unlike VR, where the entire scene is digitally rendered, AR allows participants to perceive the real world in real time while virtual objects are seamlessly integrated into their field of view [13, 14]. This offers several key advantages: (i) AR preserves the natural structure, depth cues, and dynamics of the physical environment while still permitting controlled experimental manipulation; (ii) participants can move freely and naturally within the scene without the constraints of a fixed setup or full virtual immersion; and (iii) AR tasks can be embedded into ecologically valid contexts such as navigation, search, or interaction with real surfaces. By combining real-world perception with controlled experimental probes, AR provides an attractive compromise between ecological validity and experimental control.

In our AR experiments, we deployed the experiment in Meta Quest Pro, we ask whether the classification effect observed in the previous online studies—namely, that a model can predict human correct/incorrect outcomes from background images—also holds in real environments outside the laboratory. This step is crucial for testing whether the effect is an intrinsic property of human vision in natural contexts, rather than an artefact of image-based psychophysical designs. Ultimately, AR allows us to align visual psychophysics more closely with natural behaviour and to probe how humans detect and classify visual signals in their everyday world.

### 3.1 The design of AR experiment

To test whether the classification effect from the online experiments survives in natural environments, we implemented a detection task in AR. Observers were instructed to report the presence or absence of a briefly presented visual probe. The probe was flashed for a single frame at the display refresh rate (33 ms) and appeared at irregular intervals, with inter-stimulus intervals uniformly distributed between 1 and 4 s. Immediately after the probe disappeared, participants had a 1 s response window to provide a binary judgment using the handheld Meta Quest Pro controller. They pressed one of two buttons corresponding to “*yes*” (probe present) or “*no*” (probe absent).

At the end of each response interval, participants received visual feedback indicating whether their response was correct or incorrect. The experimental pipeline is summarized in Fig. 7. Each trial consisted of four main stages: processing, probe presentation, response, and feedback. In parallel, video recordings were collected and later decoded to extract frames for machine learning analysis.

**Figure 7:**
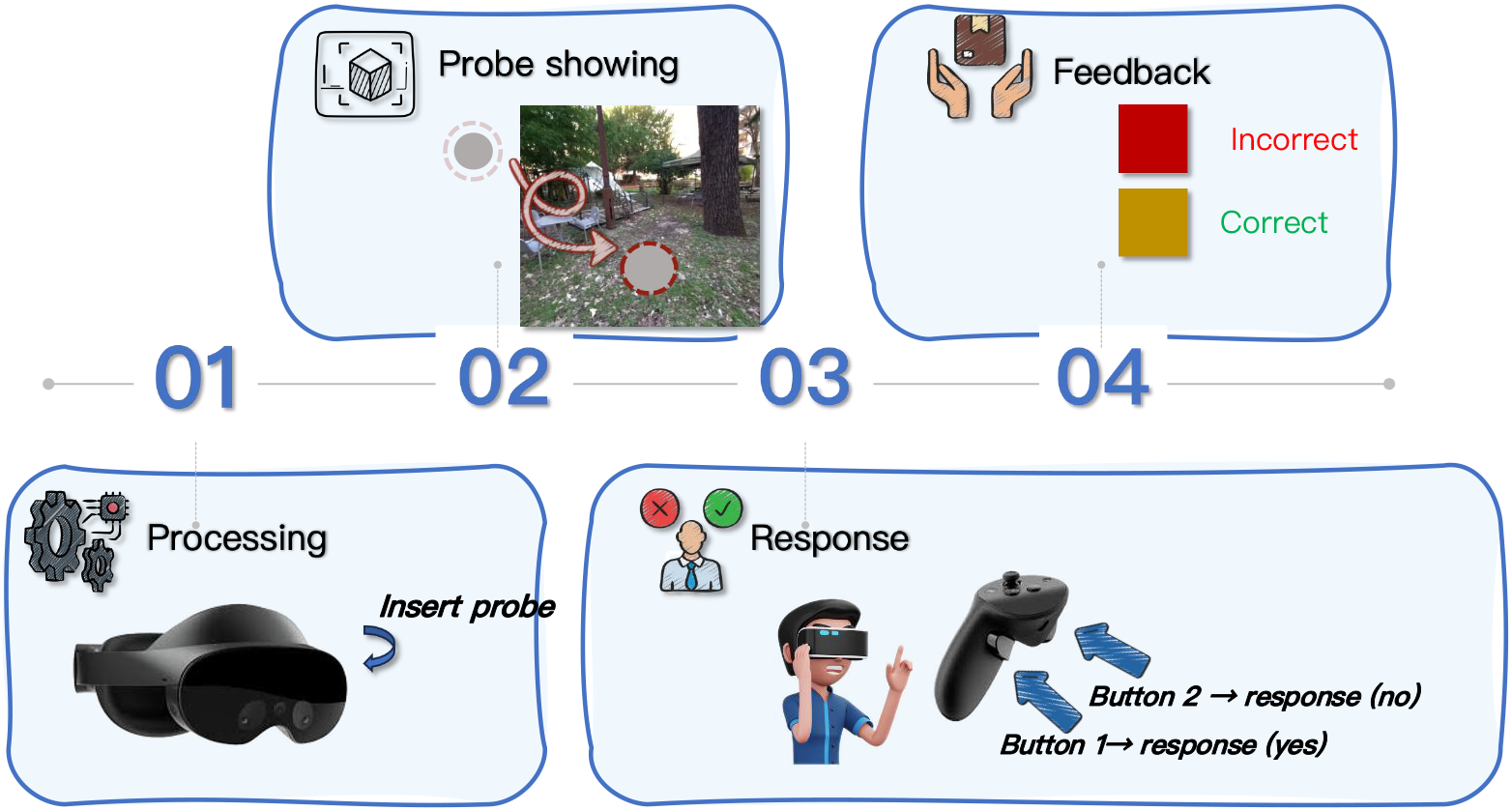
Pipeline of our AR experiment and subsequent analysis. In AR Experiment, each trial proceeds through four stages: processing, probe presentation, response, and feedback. A probe is inserted into the AR environment, the participant responds via controller button presses, and receives real-time feedback. After obtaining enough correct/incorrect trials, we use DNNs for training and testing.

In the detection task, if participants saw the probe they pressed button 1; if they did not see it, they pressed button 2. In the online experiments, we used precue, postcue, and simultaneous-cue conditions. In AR, however, participants walked within a relatively large region in the garden. If we had applied precue or postcue conditions, the participant’s movement between cue and probe would have shifted the relative position of the two, making cue–probe alignment highly unreliable. We therefore adopted the simultaneous-cue configuration from the online experiment: subjects made their decision upon seeing the cue (target-present or target-absent), within a 1 s decision window.

As in the online experiments, each trial was labeled as correct or incorrect by comparing the participant’s response to the ground truth. We downsampled correct trials to obtain equal numbers of correct and incorrect examples. These balanced datasets were then used to train DNN classifiers. The input consisted of natural images (with or without probes), and the output label was derived from human behaviour (correct versus incorrect). In this way, the model learns directly from human response data; once trained, it can take a new image as input and predict whether a human observer is likely to respond correctly or incorrectly based on the image content alone.

The task was a yes/no detection task: participants reported the absence or presence of the target bar. Magenta cues appeared at random intervals between 1 and 3 s. When the cues appeared, participants had 1 s to decide whether they saw the probe. The AR experiment thus implemented a simultaneous-cue condition: the target and cue co-occurred, and the target was visible for 33 ms in each trial.

### 3.2 YES/NO task

One example of YES/NO task is shown in Fig. 8. To better understand whether the classification effect depends on the probe itself, we designed four conditions for the AR analysis, differing in whether the probe was included or excluded from the input. Specifically, we considered four image variants for model training: *probe-containing, probe-only, probe-less*, and *preimage*, illustrated in Fig. 9.

**Figure 8:**
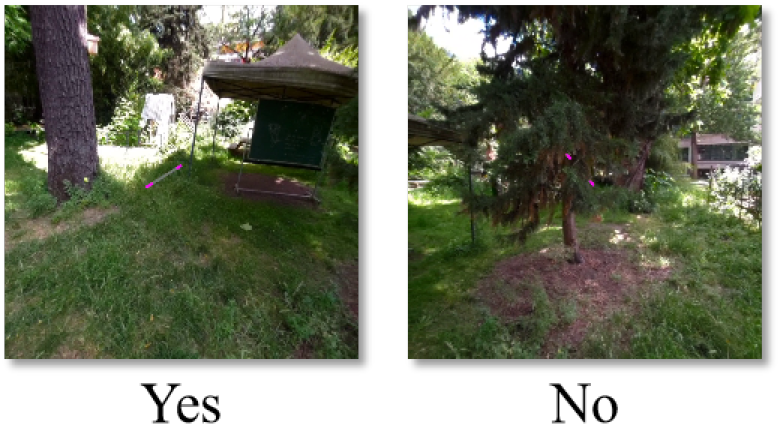
Pipeline of the YES/No task. Participants indicate whether the gray bar is presented.

**Figure 9:**
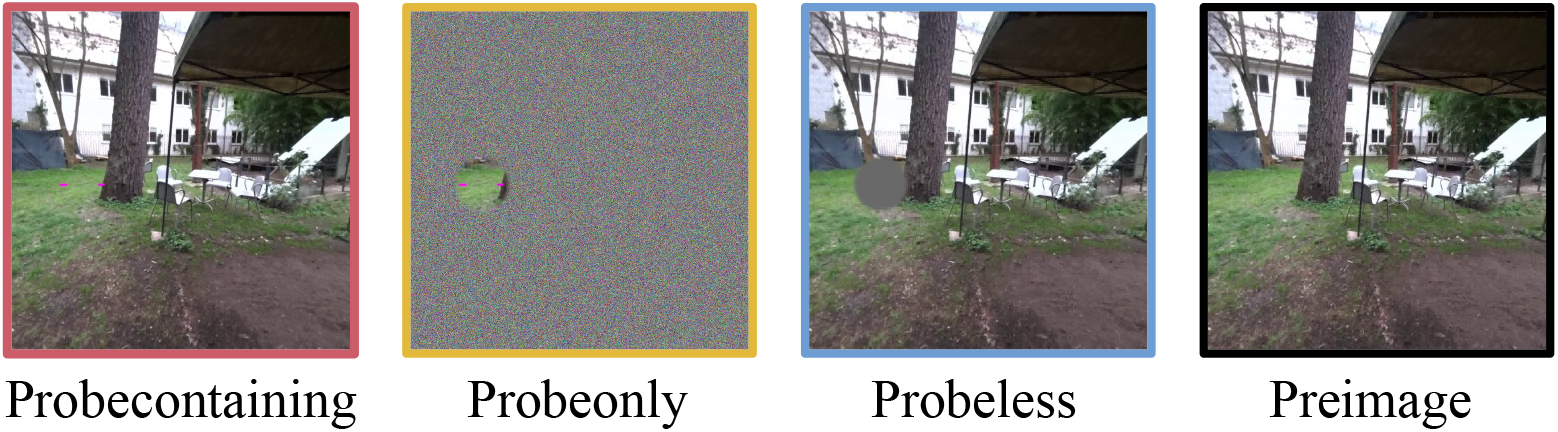
Four conditions used in the yes/no task: probe-containing, probe-only, probe-less, and preimage.

Each image had a resolution of 1024 × 1024 pixels. The cue+probe region was defined as a circular area with diameter 64 px. Video recordings were collected continuously as participants performed the AR task, and individual trials were decoded offline.

- **Probe-containing**: the exact frame seen by the participant at the moment of the probe.
- **Probe-only**: only the local region containing the probe is preserved, with the rest of the image removed. To avoid sharp edges, we applied a Gaussian filter at the patch boundary.
- **Probe-less**: the probe region is replaced by a gray disk while the rest of the image remains unchanged, again smoothed by a Gaussian filter.
- **Preimage**: a “clean” image approximating the background of the probe-containing frame. Because the cue appears for only 33 ms, we used the frame immediately preceding the probe as the preimage, which is nearly identical to the probe-containing image but without the probe.

#### 3.2.1 Participants and data preparation

We tested 10 naive participants. We ultimately used data from 8 participants, so that the same group could be used for the AR and AR-in-the-lab comparisons (see Sec. 3.4 for details). The number of usable trials per participant ranged from 554 to 1,646. In total, 10,141 trials were collected, of which 7,584 met the inclusion criteria and were used for analysis. The mean detection accuracy across included participants was 73.5%.

#### 3.2.2 Results

We trained a Swin Transformer [8] separately on the four conditions described above and evaluated performance using AUC as the primary metric. The results are shown in Fig. 10.

**Figure 10:**
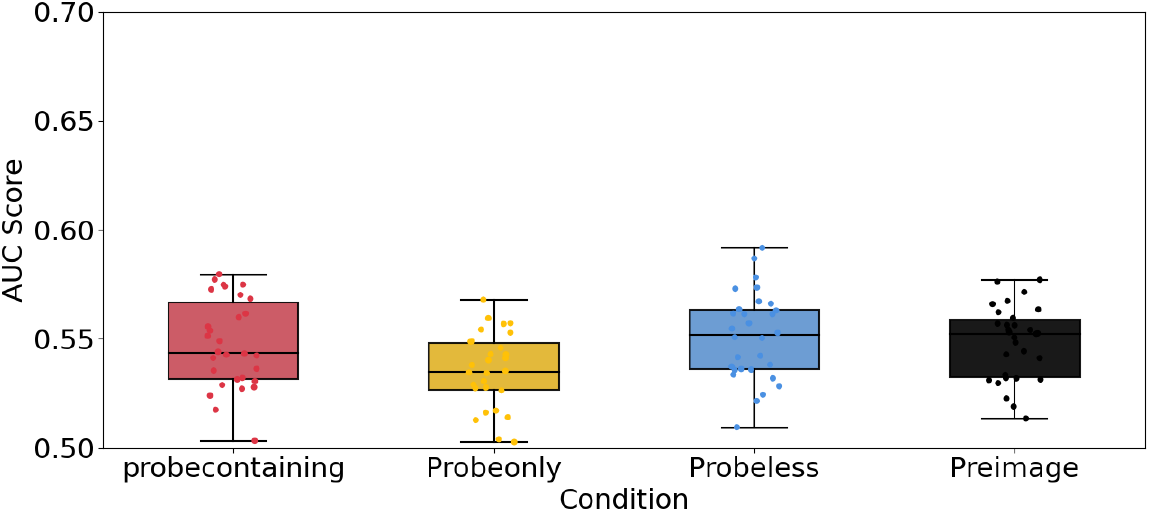
AUC for correct/incorrect prediction in the yes/no task across the four image conditions.

In all four conditions, the AUC is clearly above chance. For the *probe-only* and *probe-less* conditions, one might worry that classification is driven by spatial biases (e.g., specific probe locations). This possibility can be tested with the *preimage* condition: here, the model only sees the background frame immediately preceding the probe. The above-chance AUC in the preimage condition demonstrates that classification is not tied exclusively to the physical presence of the target but can be decoupled from it.

Conceptually, when we split trials into correct and incorrect categories, we imprint the influence of the probe onto the background images, via human behaviour. The model learns an indirect association between the preimage and the correct/incorrect label because the splitting procedure carries a “trace” of the probe through the human response. The network thus exploits systematic relationships between background structure and subsequent perceptual success or failure.

### 3.3 2AFC task

In the yes/no paradigm, participants may adopt individual response biases—some observers might respond “yes” when uncertain, others “no.” To attenuate such decision biases, we implemented a two-alternative forced-choice (2AFC) design. In the 2AFC task, two potential target locations were cued simultaneously on the left and right sides of the display, and participants reported which side contained the gray probe. The overall design is illustrated in Fig. 11, where the probe appears on either the left or the right side.

**Figure 11:**
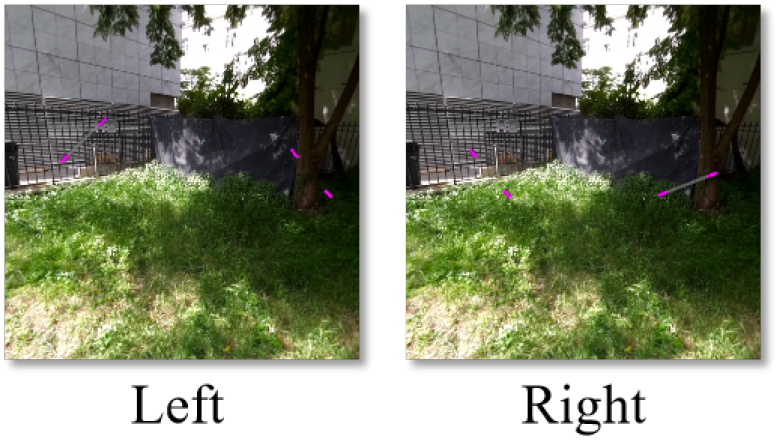
Pipeline of the 2AFC task. Two cues appear simultaneously, and participants indicate on which side (left or right) the gray bar is presented.

#### 3.3.1 Participants and data preparation

We tested nine naive participants, yielding a total of 5,101 trials for analysis, and the average final performance across included subjects was 69.6% accuracy.

#### 3.3.2 Results

Figure 12 shows the classification performance of the models trained on the 2AFC data. Using the same analytical procedures as in the yes/no task, we found that all models achieved AUC values above chance across all four conditions.

**Figure 12:**
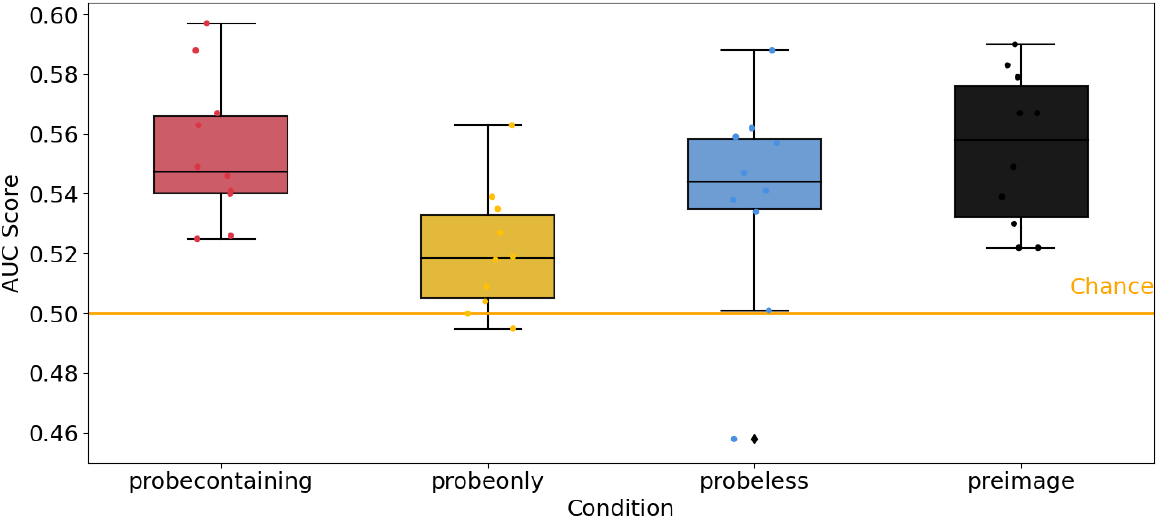
AUC for correct/incorrect prediction in the 2AFC experiment across the four image conditions.

These findings confirm that the classification effect persists even under a bias-reduced 2AFC paradigm. In the *preimage* condition, where the model receives only background images without a probe, AUC remains above chance—demonstrating that global image context still predicts whether a participant is likely to make a correct or incorrect judgment.

Together, the results from the yes/no and 2AFC paradigms show that the model–human correspondence is not an artefact of response bias or decision strategy. Instead, it reflects an intrinsic link between natural image structure and human visual detectability, which deep neural networks can partially capture and reproduce. The robustness of this effect across task structures suggests that the predictability of human visual performance by DNNs generalizes beyond a single decision framework.

### 3.4 AR in the lab

To further verify the robustness of the classification effect observed in AR, we conducted a follow-up “AR-in-the-lab” experiment as a sanity check. In this task, we re-tested the participants who had previously completed the outdoor AR yes/no experiment. All scene images (probe-present and probe-absent) that they had encountered in AR were replayed on a large display screen in the lab, approximating the visual content of the original AR sessions. Participants were asked to indicate whether a probe was present or absent in each image, thereby generating a new set of correct and incorrect labels. We then repeated the correct/incorrect classification analysis using the same computational framework as in the outdoor AR condition, but with labels derived from indoor responses.

#### 3.4.1 Participants and data preparation

Ten participants originally took part in the outdoor AR yes/no experiment; eight of them were available to repeat the task in the laboratory. These eight participants produced a total of 7,584 trials with average accuracy of 84%.

#### 3.4.2 Results

As shown in Fig. 13, when correct/incorrect labels were derived from participants’ laboratory responses, the classification effect remained clearly present. Moreover, the effect magnitude was higher indoors than outdoors, indicating that the phenomenon is not dependent on AR-specific factors. Instead, it generalizes across both naturalistic (outdoor AR) and controlled (indoor lab) viewing conditions.

**Figure 13:**
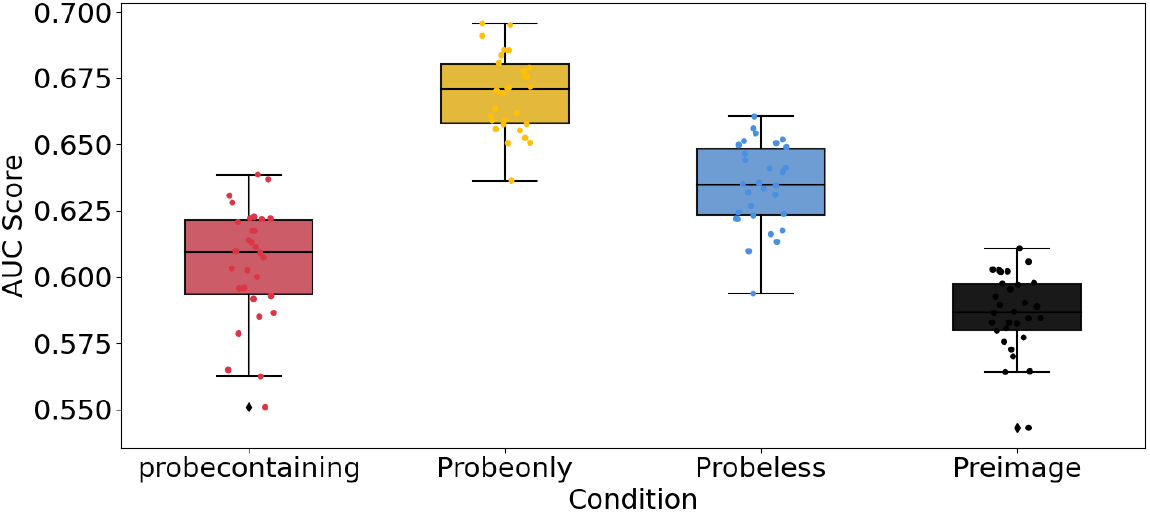
Results of the “AR-in-the-lab” experiment. The classification effect persists under laboratory conditions, with higher AUC than in the outdoor AR task.

## 4 Isolating relevant scene features via machine learning

During perceptual decision-making, the visual system must rapidly extract and evaluate task-relevant information from the environment. Contemporary computational accounts propose that evidence in favor of a choice accumulates over time until it reaches a decision threshold [15, 16]. Here, we ask how this evidence-formation process is shaped by the statistical structure of natural scenes [17]. Prior work has shown that neural responses to natural images are strongly modulated by scene complexity, suggesting that cortical processing is sensitive to—and potentially tuned for—the regularities of real-world environments [18, 19]. This motivates the central question of this chapter: *which aspects of natural scenes most strongly influence human visual decisions?*

### 4.1 Dataset

Across tasks, images were drawn either from COCO [1], SA-1B [2] or homemade AR dataset. In general, the total number of trials used in our experiments is shown in Fig.14. This section presents the results of the analysis of most tasks; the remaining “EEG image” and “Unrelated” datasets are not mentioned here.

**Figure 14:**
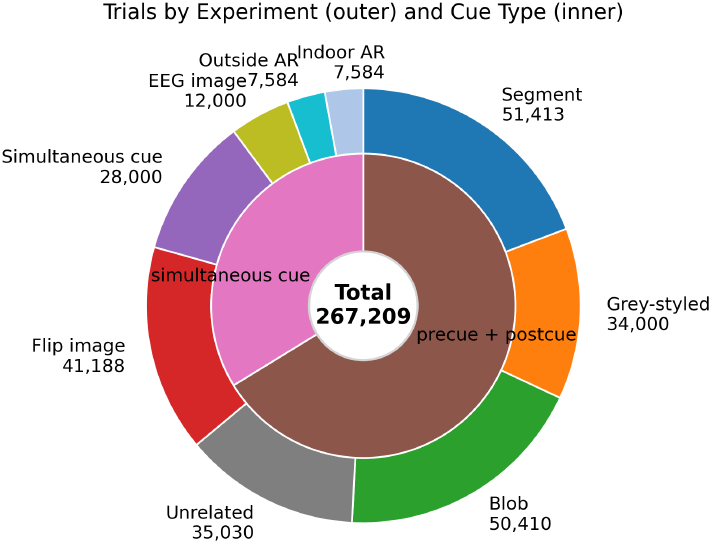
Data volume distribution across experiments. The outer ring shows the number of trials for each experimental condition, while the inner ring aggregates the data into two cue presentation types: precue+postcue (brown) and simultaneous cue (pink). The center displays the total number of trials collected (n=267,209). Color coding corresponds to individual experiments.

### 4.2 Analysis in psychophysics experiments

#### 4.2.1 Image-driven separation of correct and incorrect trials

To identify visual properties that systematically differentiate correct from incorrect outcomes, we performed a feature-based analysis centered on elementary image statistics. Across all online psychophysics tasks—including the segment, blob paradigms—the same two features consistently emerged as the most informative: **texture entropy** and **edge density**.

Figure 15 summarizes this effect across the online tasks. The model outputs both a predicted label and an associated confidence score, reflecting the certainty of each decision. To examine whether low-level image structure systematically relates to human behavioral outcomes, we selected the 100 trials with the highest classification confidence within each behavioral category (correct and incorrect). By focusing on high-confidence cases, we aimed to identify prototypical examples of stimuli that consistently led to correct or incorrect responses. When projected into the two-dimensional feature space defined by texture entropy and edge density, these high-confidence correct and incorrect trials occupied partially distinct regions, suggesting that differences in low-level structural properties may contribute to variations in human detection performance. Correct trials tend to cluster in regions characterized by lower entropy and sparser edge structure, whereas incorrect trials are associated with higher structural complexity and denser edge patterns.

**Figure 15:**
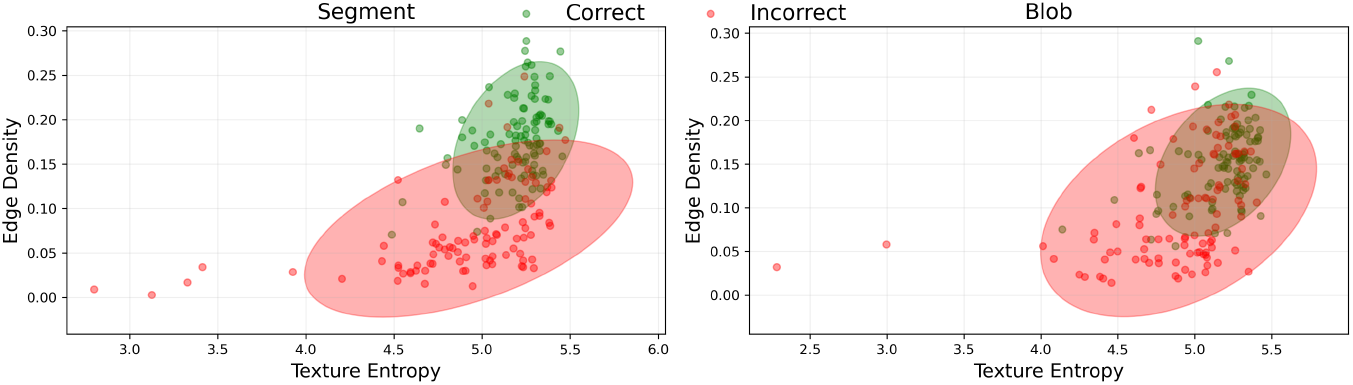
Feature-space distribution of correct and incorrect trials (postcue condition) across the online psychophysics paradigms, shown in the texture-entropy × edge-density space.

Crucially, this separation pattern is consistent across paradigms: despite changes in probe type, timing, stimulus format, or image types, correct and incorrect trials remain differentiable using these *global* image statistics. This provides converging evidence that perceptual outcomes are systematically shaped by scene-level structure, rather than by local probe properties or probe geometry.

To provide intuition for what the feature space captures, Fig. 16 shows representative images from the most discriminative regions of the space.

**Figure 16:**
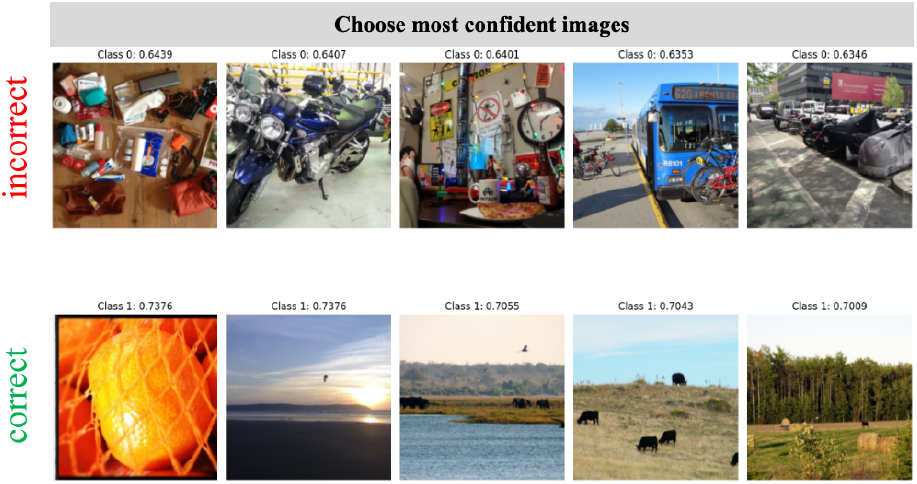
Most discriminative images with highest confidence selected by DNN. Top: incorrect trials; bottom: correct trials.

### 4.2.2 Low-dimensional models recover the same structure

We next asked whether these two global statistics are sufficient to support predictive modeling without access to pixel-level input. We trained conventional machine-learning classifiers—SVM [20], Random Forest [21], and Logistic Regression [22]—using only texture entropy and edge density as features. As shown in Fig. 17, despite operating in a two-dimensional space, these models achieve reliably above-chance performance across tasks.

**Figure 17:**
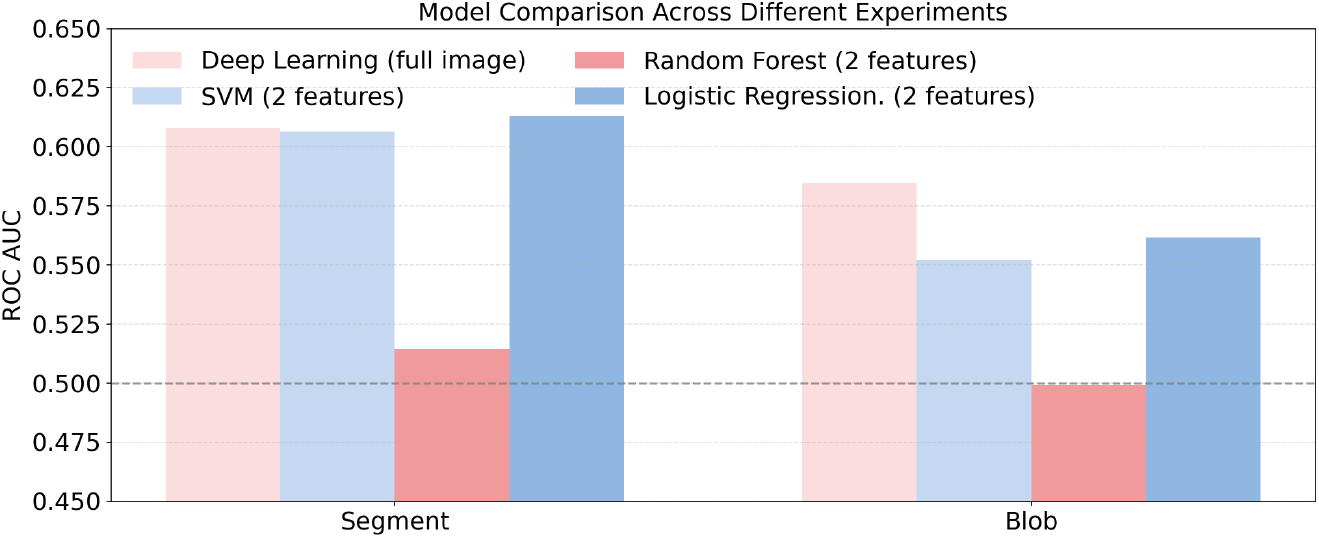
Classification performance of conventional classifiers trained only on texture entropy and edge density.

Importantly, this result mirrors the behavior of deep networks trained on full images, while making the driving factors explicit: a substantial fraction of the predictability arises from simple, interpretable *global* scene statistics rather than high-level semantics or localized probe-related features.

Taken together, the psychophysics experiments establish a robust, cross-paradigm finding: human correct and incorrect in natural-image detection tasks is strongly modulated by global scene structure. Texture entropy and edge density capture a large component of this dependency, enabling both deep and shallow models to predict correct versus incorrect outcomes from probe-free backgrounds.

### 4.3 Analysis in AR experiments

We next asked whether the image-driven effects observed in screen-based psychophysics persist under more naturalistic viewing. The AR experiments provide a critical test: perceptual decisions unfold in stable, three-dimensional environments with natural depth cues and unconstrained head/body movement, yet our computational analyses still operate on *probe-free* images to isolate the contribution of global scene context.

**Outdoor AR**. In the outdoor condition, participants performed the detection task while viewing real scenes with full depth structure and natural variability in illumination, layout, and background statistics. This setting therefore evaluates whether global scene properties continue to bias perceptual outcomes under ecologically realistic conditions.

**Indoor AR**. The indoor condition re-tested the same image set in a controlled laboratory environment, substantially reducing environmental variability (e.g., lighting fluctuations and background diversity). This provides a complementary regime in which global scene statistics are more constrained, allowing us to test when low-dimensional descriptors remain informative.

Figure 18 summarizes both the feature-space distributions and the performance of conventional classifiers in the two AR conditions.

**Figure 18:**
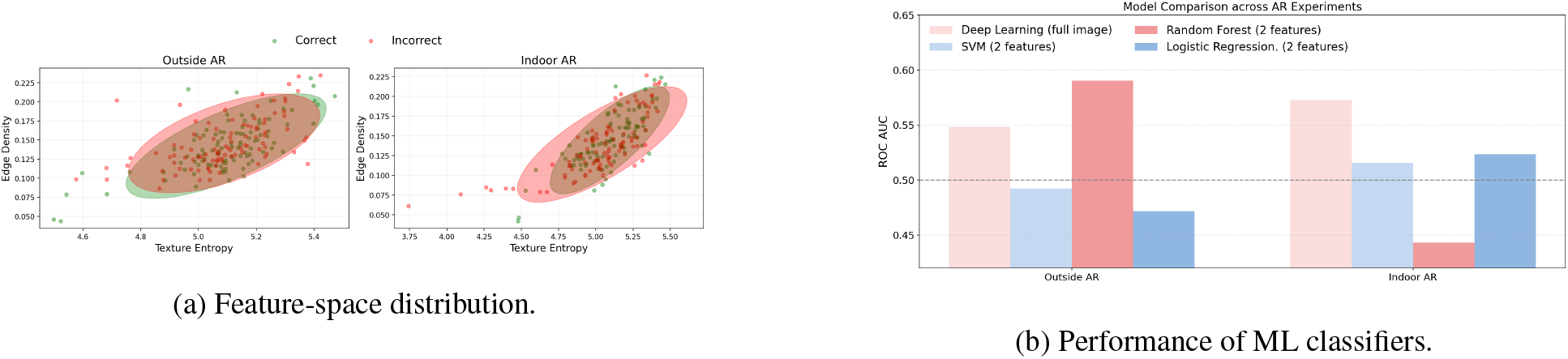
Feature-space separation (texture entropy× edge density) and conventional ML performance in the outdoor and indoor AR experiments.

In the outdoor AR condition, **texture entropy** and **edge density** provide only *partial* separation between correct and incorrect trials. Nevertheless, a Random Forest classifier trained on these two global statistics achieves reliable above-chance AUC, in fact outperforming the DNN in this setting. This pattern aligns with the psychophysics results: even under naturalistic viewing and free movement, coarse *global* scene statistics still carry predictive information about whether observers will succeed or fail on a given trial.

In the indoor AR condition, by contrast, the DNN trained on full images clearly outperforms the feature-based models. This divergence suggests that when global scene variability is strongly reduced, trial-wise perceptual outcomes may depend on more subtle, higher-order, or spatially structured cues that are not well captured by two scalar descriptors such as entropy and edge density.

Overall, the AR experiments refine the picture established by psychophysics. In richly varying environments, simple global statistics (entropy and edge density) capture a substantial component of context-driven perceptual variability. In visually constrained environments, these low-dimensional descriptors become insufficient even though global context remains predictive, indicating that the *relevant* global cues can shift with the environment.

Taken together, these results support a single, coherent conclusion across the thesis: perceptual correct and incorrect is primarily governed by *global scene context*, but the specific global cues that best explain performance depend on the statistical regime of the environment. Texture entropy and edge density provide a robust and interpretable account in diverse natural scenes, while more expressive scene descriptors are likely required when variability is limited.

## 5 EEG responses as neural signatures of image-driven perceptual outcomes

Behavioral responses alone cannot reveal how the brain represents and processes the visual information that ultimately drives correct versus incorrect judgments. To address this gap, this section investigates the neural correlates of perceptual outcomes by analyzing EEG signals recorded during the same detection tasks.

### 5.1 Experiment setting

The experimental setup is illustrated in Fig. 19. In particular, the *preimage window* (0–0.55 s) includes the full fade-in and static presentation of the natural scene, but no probe. This window is therefore ideally suited to testing whether EEG reflects image-driven differences between trials that will ultimately be correct versus incorrect, independent of any probe-related response.

**Figure 19:**
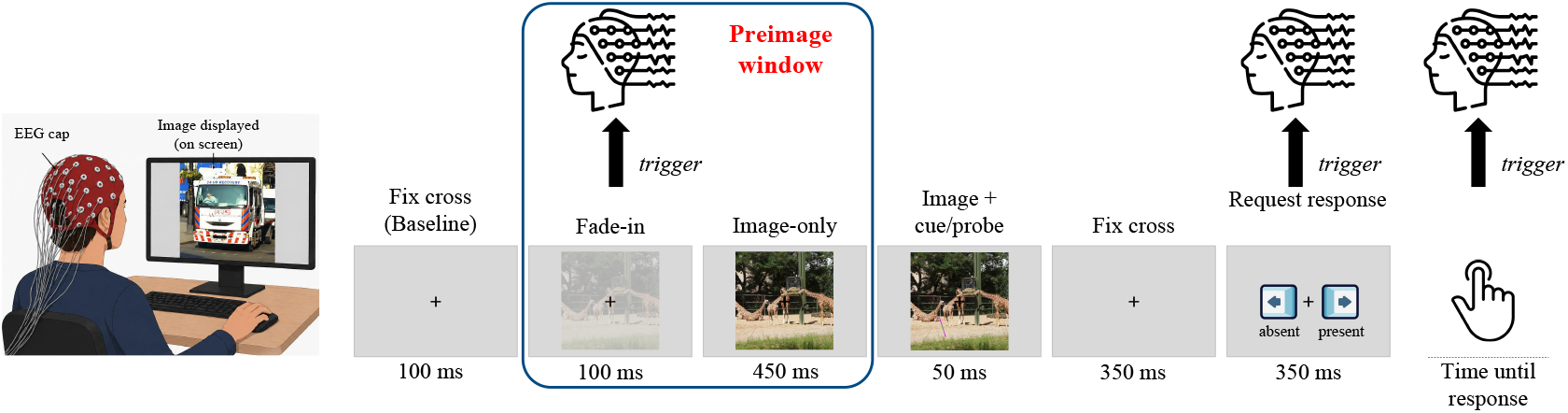
EEG experimental setup. Within the first 100 ms, the natural image fades in linearly from transparency 0 to 1. The fully visible image is then presented for 450 ms, after which the target (cue + gray segment, or cue alone) appears for 50 ms. The fixation cross remains for a further 350 ms, followed by the response period. Feedback is conveyed by the color of the cross (green for correct, red for incorrect).

### 5.2 Participants and data preparation

EEG data were recorded while participants performed the same natural-image detection task used in the online psychophysics studies. Across all sessions, we collected 12,000 trials from 8 participants. EEG data preprocessing followed these steps: data were firstly re-referenced to the average of 64 scalp electrodes, followed by a 4th-order bandpass filter (0.1–40 Hz). Ocular artifacts were removed using independent component analysis (ICA). Data were then epoched from *−*0.1 to 1 sec relative to image onset (sampling rate = 1024 Hz, step = 0.00097656 sec) and baseline-corrected by subtracting the average activity of the 100 ms pre-stimulus period. All epochs were retained in chronological order, matching the corresponding behavioral data file. To construct a balanced dataset for training, we retained 2,594 trials from each behavioral category, including both EEG recordings and their corresponding images. Each trial was labeled according to participants’ behavioral responses.

For decoding analyses, we used an 80/20 split: 80% of trials were used for model training and validation, and the remaining 20% served as a held-out test set. All decoding results reported below are averaged over 10 independent runs, and we report mean ± standard deviation of AUC and accuracy.

### 5.3 EEG Topography Visualization

To characterize the spatial distribution of discriminative information, we employed an electrode-kernel decoding approach based on frequency-domain information. For each electrode, a local kernel was constructed by selecting the center electrode together with its three nearest neighboring electrodes in Euclidean space (kernel size =3), resulting in a four-channel feature set. Decoding performance was quantified by training a classifier using only the signals from the electrodes within each kernel and computing the AUC value. The resulting AUC value was then assigned to the center electrode. Repeating this procedure for all electrodes yielded a kernel-based decoding topography, reflecting the local spatial distribution of information relevant for condition discrimination. The measured the channel distribution across the scalp is shown in Fig.20.

**Figure 20:**
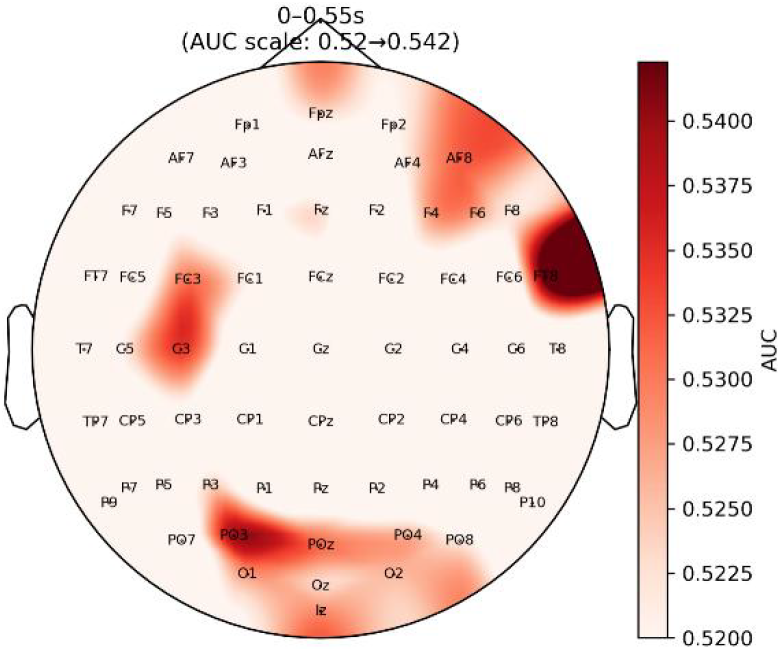
Electrode-kernel decoding topography across all canonical oscillatory bands.

By decoding from the occipital, this preliminary result show “visual” brain for the early part. The window corresponds to the background-only period before target onset. During this phase, decoding performance is strongest over fronto-central electrodes, suggesting that discriminative information predominantly reflects background-driven preparatory state differences.

### 5.4 Dual-modal fusion of EEG and image features

Having established that EEG alone carries above-chance information about perceptual outcomes in the preimage window, we next asked whether combining EEG with image-derived features improves prediction of correct vs. incorrect decisions. Crucially, we also tested whether the benefit of fusion depends on trial-by-trial alignment between EEG and image, which would indicate that neural responses are specifically tuned to the particular scene shown on each trial rather than merely reflecting generic difficulty or decision bias. We employed two fusion ways, i.e., trial-by-trial match and label-match. The contrast between the two conditions is illustrated in Fig. 21. If EEG and image features encode complementary, image-specific information, fusion performance should be highest when trial identity is preserved (trial-by-trial match) and progressively degrade toward chance as trial-level alignment is disrupted (shuffle conditions). The label-match condition serves as a critical control, testing whether any fusion benefit can be explained solely by generic correct/incorrect signals—such as overall task difficulty—rather than by trial-specific cross-modal correspondence.

**Figure 21:**
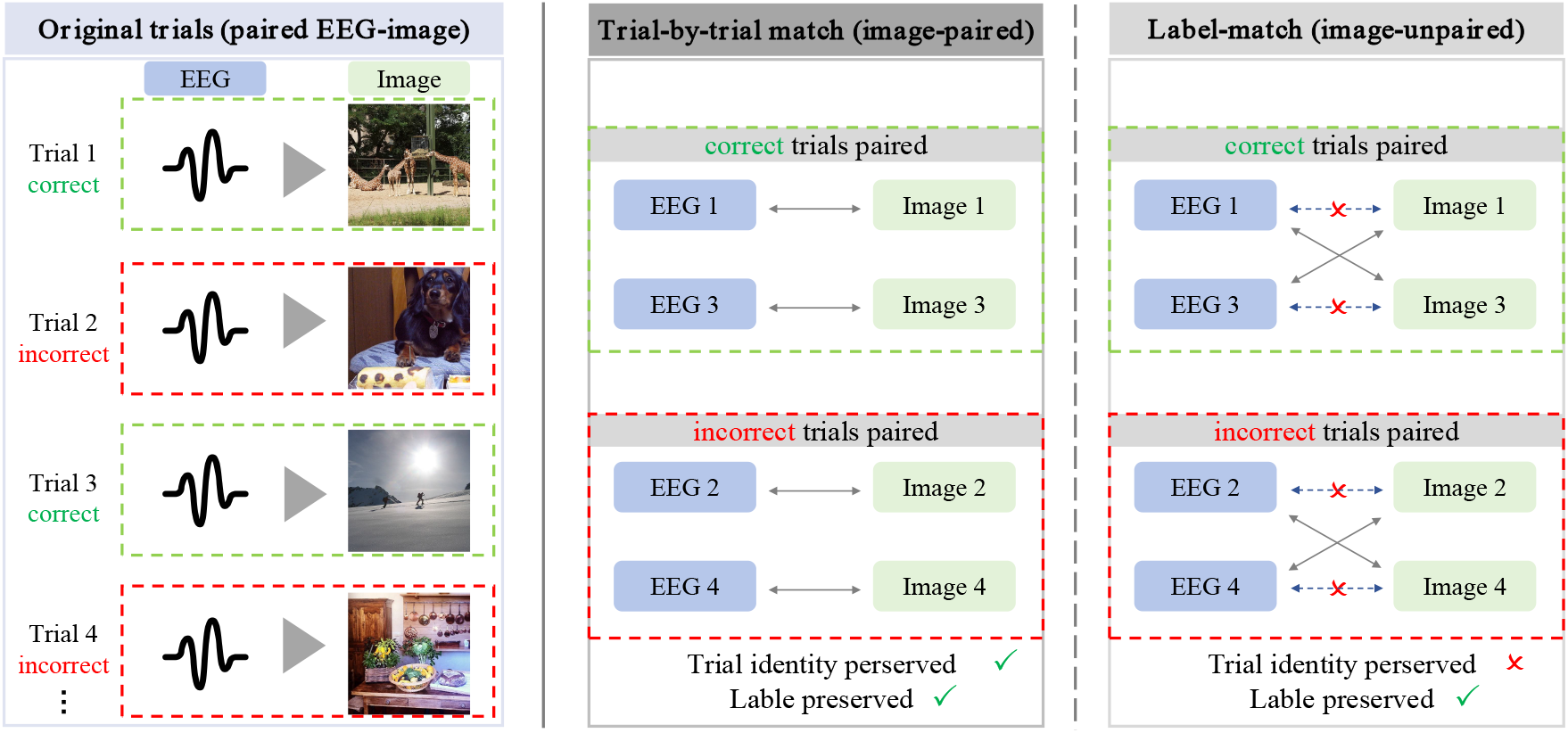
Fusion conditions. *Left*: Original EEG experiment procedure, where each EEG signal corresponds to a specific image on a given trial. *Middle*: Trial-by-trial match, in which EEG and image features from the same trial are paired. *Right*: Label-match, where EEG and image features come from different trials but share the same correct/incorrect label.

For the preimage window, we use frequency-domain EEG features, reflecting early visual encoding, and fuse them with image-based features from the corresponding natural scenes. Fig. 22 summarize the results. The AUC values were obtained from 10 independent model training runs, and the error bars represent the std across these runs.

**Figure 22:**
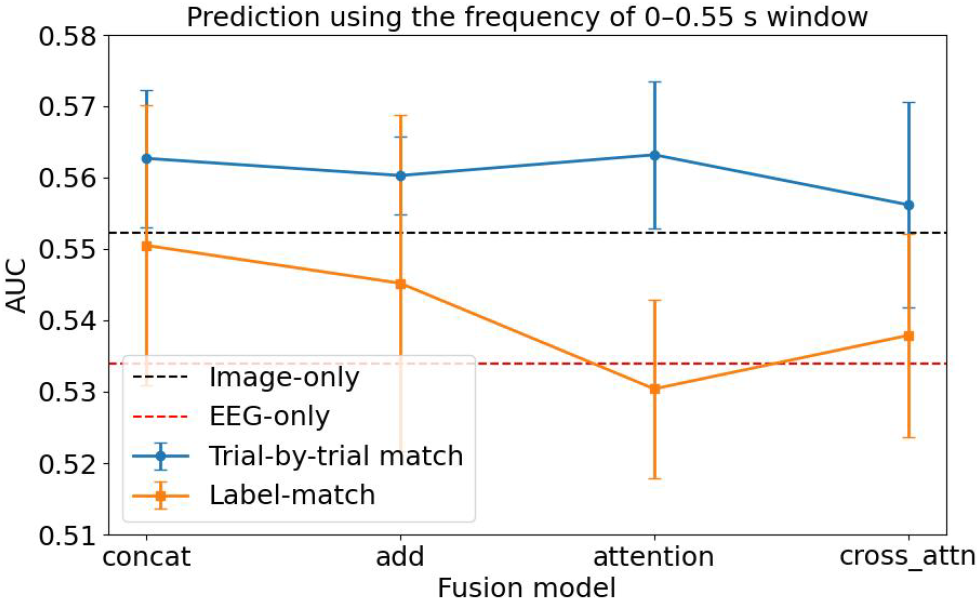
Performance of different fusion and control conditions for the probe-free window (frequency-domain EEG). The x axis shows different fusion methods, i.e., “Concatenation”, “Add”, “Attention” and “Cross-attention”.

Two main observations emerge:

- **Fusion improves decoding**. For all four fusion schemes, trial-by-trial fusion yields AUCs above 0.56, higher than both EEG-only (AUC≈ 0.53) and image-only (AUC ≈0.55) baselines. This confirms that EEG and image features carry complementary information about perceptual outcomes.
- **Trial specificity matters**. Trial-by-trial fusion consistently outperforms label-match fusion, where EEG and images share the same correct/incorrect label but come from different trials.

Thus, even in the preimage window—before the probe appears—EEG contains scene-specific information that, when combined with DNN-derived image features, enhances prediction of whether a given trial will be correct or incorrect.

In this section, we used EEG to examine whether neural activity carries information about image-driven perceptual outcomes in near-threshold detection tasks. Three main findings emerge. First, EEG information showed a temporal dissociation: in the *preimage window* (0–0.55 s), when no probe was visible, frequency-domain features yielded above-chance decoding whereas time-domain features did not; Second, combining EEG with DNN-derived image features improved prediction relative to either modality alone, indicating that neural and image-based representations provide complementary information about trial outcomes. Third, this fusion benefit was strongest when EEG and image features were paired at the trial level. Trial-by-trial fusion outperformed label-match fusion and shuffle controls, demonstrating that early EEG signals are linked to the specific scene content of each trial rather than to generic correct/incorrect labels alone.

Together, these results suggest that even before the probe appears, EEG contains scene-specific information that complements image-based features and helps predict whether a given trial will be perceived correctly.

## 6 Conclusion

Understanding how humans extract task-relevant information from rich natural scenes remains a central challenge in vision science. In this study, we combined controlled psychophysical experiments, AR-based naturalistic viewing, computational image analysis, deep neural networks, and EEG measurements to examine how background scene structure shapes perceptual decision-making in near-threshold detection tasks. Across these converging approaches, we found that perceptual accuracy is not determined solely by the target itself, but is systematically modulated by the statistical structure of the surrounding scene.

The online psychophysical experiments first showed that human detection performance varies reliably across background images, even when the probe stimulus is held constant. This effect was especially pronounced in postcue conditions, where observers had less opportunity to deploy anticipatory spatial attention and therefore relied more strongly on information available from the scene. Deep neural networks trained only on probe-free background images were able to predict whether observers would respond correctly or incorrectly, indicating that natural scenes contain reproducible statistical cues that bias perceptual outcomes.

The AR experiments extended this finding to more naturalistic viewing conditions. When probes were embedded in real-world environments viewed through a head-mounted display, observers experienced natural head and body movements, variable illumination, and richer environmental context. Despite this additional complexity, probe-free background information still predicted trial-level performance. This suggests that context-driven modulation of perception is not an artifact of computer-screen experiments, but a robust feature of natural visual behavior.

To identify the image properties underlying these effects, we analyzed both low-level statistics and model-internal representations. Texture entropy and edge density emerged as particularly informative features: scenes with higher entropy and denser edge structure were associated with lower detection performance, and these features alone supported above-chance classification of correct versus incorrect trials. These results provide a concrete account of how low-level scene statistics can influence perceptual variability, independently of high-level object semantics.

Finally, EEG analyses showed that scene-driven differences in perceptual outcome are reflected in neural activity. In the preimage window, when the natural scene was visible but the probe had not yet appeared, frequency-domain EEG features contained information predictive of later trial outcome. Moreover, combining EEG features with DNN-derived image representations improved decoding performance beyond either modality alone. Trial-by-trial fusion outperformed label-matched and shuffled controls, suggesting that the neural signal was linked to the specific scene content of each trial rather than to generic correct/incorrect labels alone.

Together, these findings support a unified conclusion: natural scene statistics systematically shape perceptual decisions before and during target detection. Background structure provides behaviorally relevant information, computational models can decode this information from probe-free images, and neural activity reflects sensitivity to these scene-specific properties. By linking psychophysics, AR, image-based modeling, and EEG, this work demonstrates that perceptual success in natural scenes depends on the interaction between external scene statistics and internal neural representations.

## Notes

### Competing Interest Statement

The authors have declared no competing interest.

